# A novel role for SHARPIN in amyloid-β phagocytosis and inflammation by peripheral blood-derived macrophages in Alzheimer’s disease

**DOI:** 10.1101/732164

**Authors:** Dhanya Krishnan, Ramsekhar N Menon, Mathuranath PS, Srinivas Gopala

## Abstract

**INTRODUCTION:** Defective immune cell-mediated clearance of amyloid-beta (Aβ) and Aβ-associated inflammatory activation of immune cells are key contributors of Aβ accumulation and neurodegeneration in Alzheimer’s disease (AD), however, the underlying mechanisms remain elusive.

**METHODS:** Differentiated THP-1 cells treated with Aβ and AD patient-derived macrophages were used as *in-vitro* model. The role of SHARPIN was analysed in differentiated THP-1 cells using siRNA-mediated knockdown followed by immunoblotting, ELISA, real-time PCR, immunoprecipitation and flow cytometry. Differentiated SHSY5Y cells were used to study inflammation-mediated apoptosis.

**RESULTS:** SHARPIN was found to regulate Aβ-phagocytosis and NLRP3 expression in THP-1 derived macrophages. Further, it was found to promote macrophage polarization to an M1 (pro-inflammatory) phenotype resulting in enhanced inflammation and associated neuronal death, demonstrated using *in-vitro* culture systems. SHARPIN expression by blood-derived macrophages was further found to be higher in the early stages of AD, which correlates with Aβ_40/42_ concentration in the plasma and age of the study subjects.

**DISCUSSION:** The novel protein, SHARPIN has been shown to play critical roles in regulation of Aβ-phagocytosis and inflammation in AD and the mechanism by which SHARPIN is activated by Aβ in macrophages has been elucidated.

## 1. Introduction

Alzheimer’s disease (AD) is pathologically characterized by the accumulation of amyloid-beta (Aβ) and hyperphosphorylated tau proteins [1]. During normal physiology, Aβ level in the brain is maintained through a homeostasis in production and degradation [2]. This is effectuated mainly by immune cells, namely microglia in the brain and macrophages and monocytes in the peripheral system through Aβ phagocytosis and degradation [3, 4]. Under certain conditions mostly associated with aging, these cells fail to phagocytose Aβ, leading to a dysregulation in the balance between Aβ production and degradation [5, 6]. In the long term, excessive Aβ accumulation accompanied by reduced degradation leads to chronic inflammatory activation of the immune cells causing neuronal death resulting in the progression of AD [7, 8]. However, mechanisms that underlie inefficient Aβ phagocytosis and enhanced inflammation by macrophages remain insufficiently addressed.

The NLRP3 (NOD-like receptor protein 3) inflammasome, a protein complex composed of NLRP3, the adaptor protein Apoptosis-associated Speck-like protein containing a CARD (ASC) and the inflammatory caspase-1, is responsible for the cleavage and maturation of inflammatory cytokines like IL-1β and IL-18 [9]. Although NLRP3 has been linked to the progression of AD [10, 11], studies focussing on the regulatory mechanism of the protein in AD have not been reported. SHARPIN (SHANK-associated RH domain-interacting protein), a part of the LUBAC (linear ubiquitination assembly complex) has been proven to be controlling the expression of NLRP3 through NF-κB activation [12]. Nevertheless, its functional role in AD has not been studied yet.

In the present study, we sought to address the role of SHARPIN and its impact on macrophage function in AD scenario. To this end, we present an evidence for the critical role of Aβ-induced SHARPIN in Aβ phagocytosis, NLRP3 expression and macrophage polarization to M1 (pro-inflammatory) phenotype in differentiated THP-1 cell line as an *in-vitro* model and SHARPIN silencing protects neurons from Aβ-induced inflammatory damage using differentiated SHSY5Y cell line. Further, utilizing control, mild cognitively impaired (MCI) and AD patient-derived macrophages, we show that SHARPIN plays a significant role the progression of AD.

## 2. Materials and methods

### 2.1. Inclusion of study subjects

AD and MCI patients were recruited from the Memory & Neurobehavioral Clinic (MNC) at the Sree Chitra Tirunal Institute for Medical Sciences and Technology (SCTIMST), Kerala, after obtaining Institutional Ethical Clearance. All the recruited subjects were tested for hypertension, hyperlipidaemia, hypercholesterolemia, Vitamin B12 deficiency, thyroid dysfunction, diabetes, cardiopathy, cranial trauma or other neurological disorders. Subjects with high plasma CRP level were excluded to avoid the possibility of systemic inflammation-mediated alteration of protein expression. The diagnostic criteria of NINCDS–ADRDA [13] were used to confirm AD and MCI pathology. The severity of AD was determined using the Clinical Dementia Rating Scale [14]. The preclinical AD cases were classified as MCI using Petersen’s criteria [15], if they scored between 1.5 and 2.0 standard deviations below the corresponding age group’s population mean score on the memory tests [16] on the Addenbrook’s Cognitive Examination [17] administered in their native language of Malayalam. The study population comprised of 63 individuals with 31 AD, 13 MCI and 19 cognitively unimpaired control subjects.

### 2.2. Isolation of monocytes from blood samples

Anti-coagulated blood was layered on Ficoll-Paque (Sigma Aldrich, St. Louis, MO, USA) medium and centrifuged at 2000 rpm for 20 min. PBMCs obtained were cultured for 14 days until complete differentiation into macrophages in RPMI-1640 medium supplemented with 10% autologous serum. Autologous serum was isolated by centrifugation of coagulated blood at 2500 rpm for 15 min, complement inactivated and filtered through 0.22 μm filter.

### 2.3. Cell culture and differentiation

THP-1 cells (obtained from National Centre for Cell Sciences, Pune) were cultured in RPMI-1640 medium supplemented with 10% FBS and were differentiated into macrophages by incubating with 100 nM phorbol 12-myristate 13-acetate (PMA) for 48 h (Suppl. Fig.1S).

SHSY5Y cells (obtained from CSIR-National Institute for Interdisciplinary Science and Technology (CSIR-NIIST), Trivandrum) were cultured in RPMI-1640 medium supplemented with 10% FBS. The cells were differentiated into mature neurons using 10 μM retinoic acid (RA) in RPMI-1640 supplemented with 1% FBS for 3-4 days.

### 2.4. Aβ preparation

Lyophilized Aβ_1-42_ and HiLyte Flour 488-labeled Aβ_1-42_ (FITC-Aβ) (Abcam, Cambridge, UK and Anaspec Fremont, California, USA respectively) were reconstituted in 1% NH_4_OH to 2 mg/ml concentration and was further diluted to 1 mg/ml in 1X PBS. Aβ thus prepared was analysed using western blotting and confirmed that Aβ prepared was in the cytotoxic soluble oligomeric form (Fig.1a).

**Fig.1:**
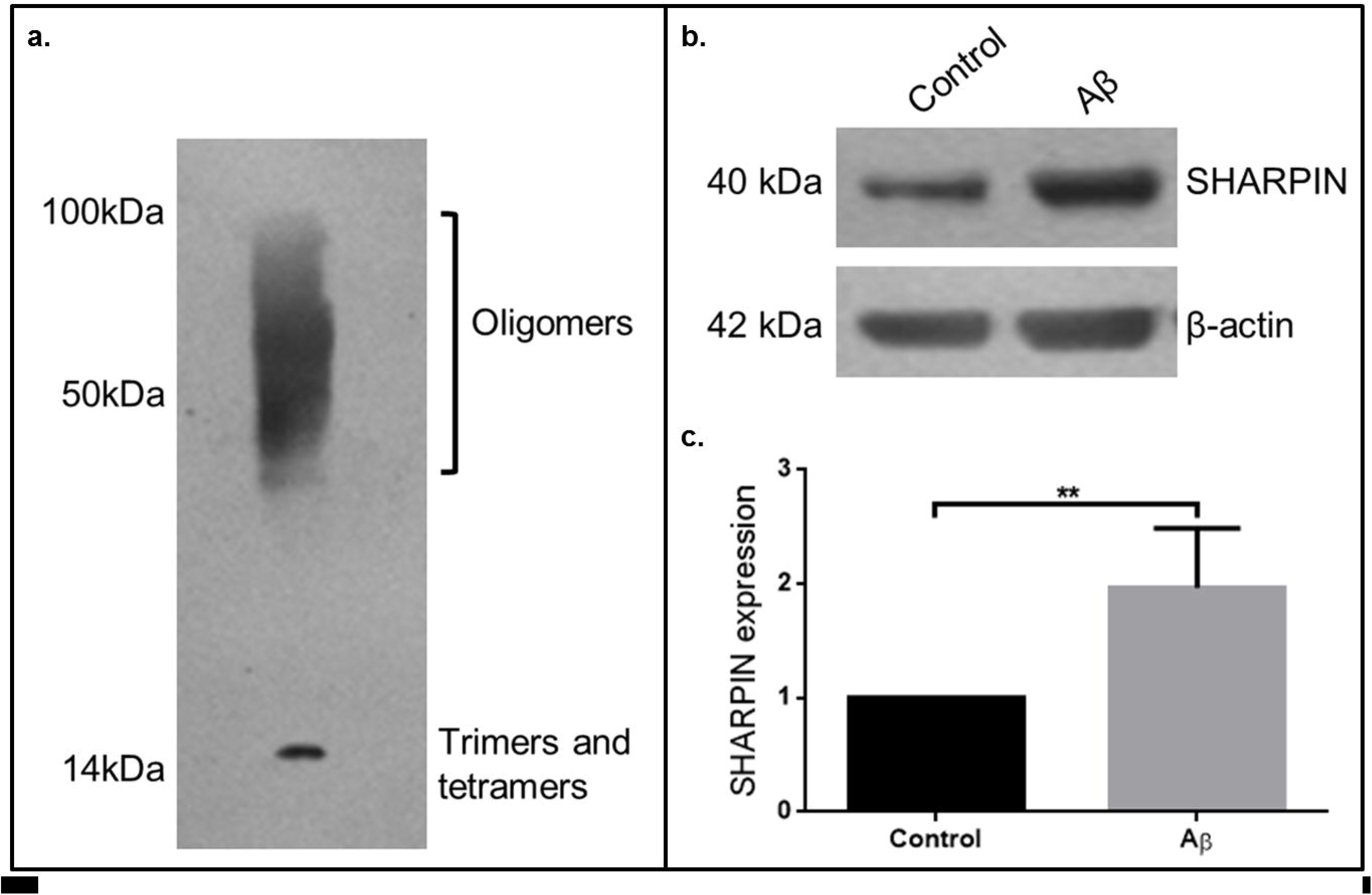
Aβ enhances SHARPIN expression. Lyophilized Aβ_1-42_ was dissolved in 1% NH_4_OH to 2mg/ml and then reconstituted in 1X PBS to 1mg/ml. The prepared Aβ was analysed using western blotting and confirmed that the Aβ prepared was in the cytotoxic soluble (oligomeric, tetrameric and trimeric) form (**a**). Differentiated THP-1 macrophages were treated with 10μM Aβ for 6h and SHARPIN expression was analysed using western blotting (**b**), quantified, normalised with control and represented graphically (Fold change in expression: Control vs. Aβ is 1: 1.97, n=4) (**c**). β-actin was used as endogenous control. Statistical analysis-student’s *t-test* with *p>0.05, **p>0.01, ***p>0.001.

### 2.5. siRNA transfection

Differentiated THP-1 cells were transfected with siRNA (Cell Signaling Technology, Danvers, Massachusetts, USA) using jetPrime PolyPlus transfection reagent (Thermofischer Scientific, Waltham, Massachusetts, USA) as per the protocol. Transfection efficiency was confirmed using western blotting.

### 2.6. Assessment of reactive oxygen species production

H_2_DCFDA (dichlorofluorescin diacetate) assay was used to detect Aβ-induced oxidative stress. The intracellular Reactive Oxygen Species (ROS) levels were quantified after pre-incubation with 10 mM N-acetyl cysteine (NAC), an ROS scavenger for 1 h and 40 μM Aβ for 12 h. The cells were trypsinized and incubated with 10 μM H_2_DCFDA for 1 h at 37°C, washed and subjected to FACS analysis.

### 2.7. Immunoprecipitation

Total protein was isolated using low-ionic isolation buffer and was incubated with primary antibody and Protein-A coated magnetic beads and pulled down by applying magnetic field. The protein bound with magnetic beads were washed, incubated with 3X Laemelli buffer at 70°C for 5 min and the purified protein was pulled down applying magnetic field.

### 2.8. Immunoblot analysis

Total protein was isolated using 1X RIPA buffer containing protease and phosphatase inhibitors and quantified using BCA assay. Denatured protein was loaded on SDS-PAGE gels for separation and were transferred to PVDF membrane, blocked with 5% skimmed milk and incubated with respective antibodies in 3% bovine serum albumin at 4°C. After overnight incubation, the washed blots were incubated with HRP-labelled secondary antibody and developed using enhanced chemiluminescence. The relative protein expression was quantified densitometrically using ImageJ software and normalized with β-actin.

### 2.9. mRNA expression analysis

Total RNA was isolated from differentiated THP-1 cells using the kit protocol (Invitrogen, Carlsbad, California, USA) and cDNA was synthesised from the isolated RNA. The mRNA expression was analysed using TaqMan Primers with human tubulin as internal control.

### 2.10. Analysis of cytokine release

The release of inflammatory cytokines and Aβ_40_, Aβ_42_ in plasma samples and THP-1 cell conditioned media were analysed using Enzyme-Linked Immuno-Sorbent Assay. The samples were pre-treated as per the kit protocol (G-Biosciences, St.Louis, Missouri, USA) and incubated in specific antibody pre-coated wells. The wells were washed and incubated with primary antibody, HRP-conjugated secondary antibody and TMB substrate sequentially and absorbance was read at 450 nm and concentration was calculated from standard plot.

### 2.11. Macrophage Aβ internalization assay

Macrophages were incubated overnight with 1μg/ml FITC-Aβ, washed with 1X PBS and examined by fluorescence microscopy. Mean Fluorescent Intensity over three different fields per sample were analysed using ImageJ software and calculated by taking the ratio of fluorescent intensity to the total number of cells in each field.

### 2.12. Conditioned media experiments

Differentiated SHSY5Y cells were treated with the conditioned media from differentiated THP-1 treated with Aβ with/without SHARPIN siRNA for 48 h and the expression of apoptotic markers were analysed.

### 2.13. Statistical analyses

One way ANOVA and student’s *t-*test were used to compare control parameters with treatment groups. Pearson’s correlation coefficient was used to correlate each parameter with SHARPIN expression in AD, MCI and control subjects. Results were represented as mean ± SEM and p value<0.05 was considered as statistically significant.

## 3. Results

### 3.1. SHARPIN regulates Aβ Receptor Expression and Phagocytosis by Macrophages

Aβ enhanced the expression of SHARPIN by approximately two-fold in THP-1 derived macrophages (Fig.1b, c). Further, we observed that knockdown of SHARPIN in macrophages significantly reduced the Aβ phagocytic efficiency using FITC-Aβ internalization assay, demonstrating a role for SHARPIN in modulating macrophage function in the context of AD (Fig. 2b, d).

**Fig.2:**
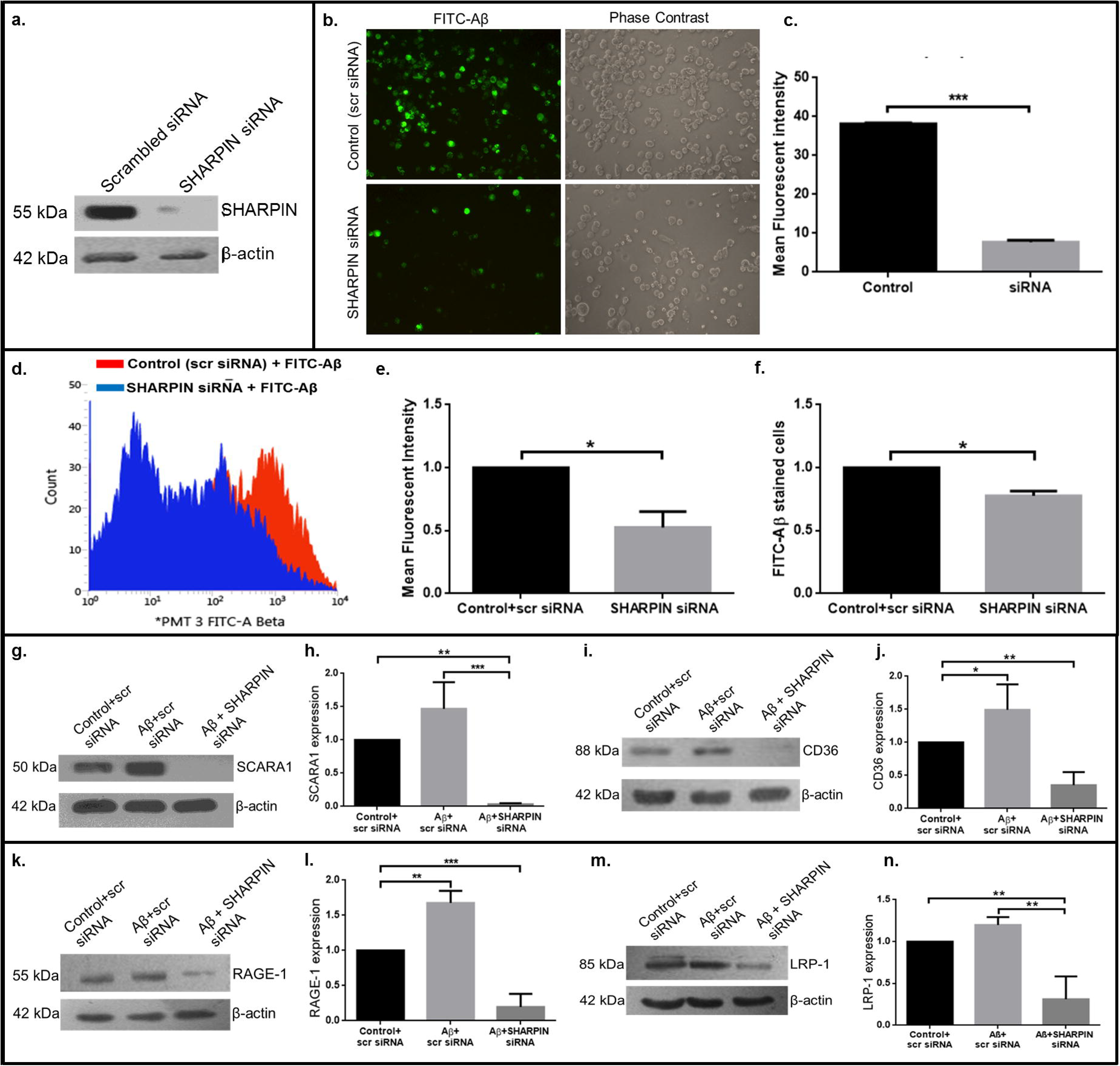
Aβ-induced SHARPIN expression regulates Aβ phagocytosis. Differentiated THP-1 macrophages were transfected with SHARPIN or scrambled (scr) siRNA and the transfection efficiency was analysed using western blotting (**a**). The efficiency of Aβ phagocytosis was analysed after incubating the transfected cells with FITC-Aβ for 12 h. Aβ uptake was analysed using fluorescent imaging (**b**) and the mean fluorescent intensity (MFI) was quantified and represented graphically (Fold change in expression: Control+scr siRNA vs. SHARPIN siRNA is 1: 1.38, n=3) (**c**). The cells were further subjected to flow cytometry analysis (**d**) to quantify fluorescence compared to the control (scr siRNA) and the MFI (Fold change in expression: Control vs. SHARPIN siRNA is 1: 1.90, n=3) (**e**) and the number of FITC-Aβ phagocytosed cells (Fold change in expression: Control+scr siRNA vs. SHARPIN siRNA is 1: 1.29, n=3) (**f**) were quantified, normalised with control and represented graphically. Differentiated THP-1 cells were transfected with SHARPIN or scr siRNA and the expression of Aβ-phagocytic receptors, SCARA1 and CD-36 (**g, i**) and receptors involved in receptor-mediated uptake of Aβ, RAGE-1 and LRP-1 (**k, m**) were analysed using western blotting, after incubating the cells with 10μM Aβ for 12 h and represented graphically after normalising with control (Fold change in expression: Control+scr siRNA vs. Aβ+scr siRNA vs. Aβ+SHARPIN siRNA is 1: 1.47: 0.033, n=4 for SCARA1; Control+scr siRNA vs. Aβ+scr siRNA vs. Aβ+SHARPIN siRNA is 1: 1.49: 0.35, n=5 for CD-36; Control+scr siRNA vs. Aβ+scr siRNA vs. Aβ+SHARPIN siRNA is 1: 1.68: 0.197, n=3 for RAGE-1; Control+scr siRNA vs. Aβ+scr siRNA vs. Aβ+SHARPIN siRNA is 1: 1.20: 0.31, n=3 for LRP-1) (**h, j and l, n**). Statistical analysis-One-way NOVA with *p>0.05, **p>0.01, ***p>0.001.

As Aβ phagocytosis is mediated by phagocytic receptors, the expression of receptors involved in Aβ phagocytosis and uptake were analyzed. SHARPIN knockdown attenuated Aβ -induced expression of the phagocytic receptors – Scavenger Receptor Class A1 (SCARA1), CD36, receptor for advanced glycation endproducts (RAGE-1) and low density lipoprotein receptor-related protein 1 (LRP-1) (Fig.2).

### 3.2. SHARPIN regulates Aβ-induced Macrophage Polarization

We showed that Aβ-induced NLRP3 expression was abolished by silencing SHARPIN in macrophages (Fig.3b), pointing to the possibility that SHARPIN and NLRP3 act in tandem to promote pro-inflammatory signaling. Further, we observed that SHARPIN knockdown results in a decrease in M1 markers (Fig.3, Suppl. Fig.3S) and increase in M2 markers (Fig.3, Suppl. Fig.3S), suggesting that Aβ-induced SHARPIN expression polarizes macrophages to pro-inflammatory M1 phenotype and SHARPIN knockdown reverses the polarization to M2 anti-inflammatory phenotype.

**Fig.3:**
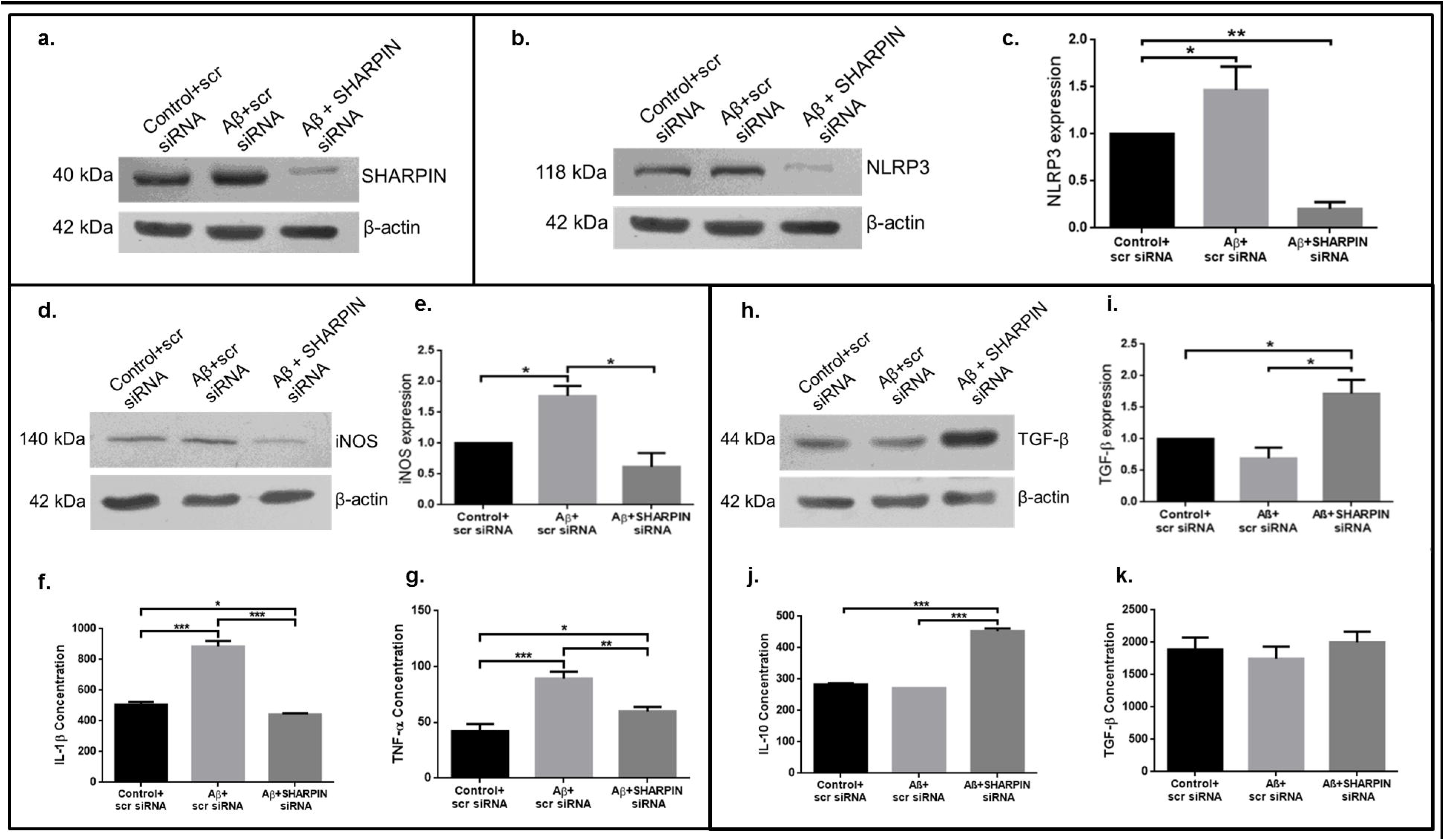
SHARPIN knockdown inhibits Aβ-mediated M1 polarization of macrophages. Differentiated THP-1 macrophages were transfected with SHARPIN or scrambled (scr) siRNA and the transfection efficiency was analysed using western blotting (**a**). The cells with/without SHARPIN knockdown were treated with 10μM Aβ for 6 h and the expression of NLRP3 was analysed using western blotting (**b**), normalised with control and represented graphically (Fold change in expression: Control+scr siRNA vs. Aβ+scr siRNA vs. Aβ+SHARPIN siRNA is 1: 1.47: 0.20, n=3) (**c**). The cells with/without SHARPIN knockdown were treated with 10μM Aβ for 12 h. The expression of M1 marker iNOS (induced Nitric Oxide Synthase) (**d, e**) and the release of pro-inflammatory cytokines (IL-1β and TNF-α) in the cell-conditioned media were quantified using ELISA and represented graphically (Fold change in expression: Control+scr siRNA vs. Aβ+scr siRNA vs. Aβ+SHARPIN siRNA is 1: 1.77: 0.62, n=3 for iNOS; Control+scr siRNA vs. Aβ+scr siRNA vs. Aβ+SHARPIN siRNA is 1: 1.75: 0.87, n=3 for IL-1β; Control+scr siRNA vs. Aβ+scr siRNA vs. Aβ+SHARPIN siRNA is 1: 2.12: 1.42, n=4 for TNF-α) (**f, g**). **D.** The expression of M2 marker TGF-β (**h, i**) and the release of anti-inflammatory cytokines (IL-10 and TGF-β) in the cell-conditioned media were quantified using ELISA and represented graphically (Fold change in expression: Control+scr siRNA vs. Aβ+scr siRNA vs. Aβ+SHARPIN siRNA is 1: 0.70: 1.72, n=3 for TGF-β; Control+scr siRNA vs. Aβ+scr siRNA vs. Aβ+SHARPIN siRNA is 1: 0.92: 1.06, n=3 for IL-10; Control+scr siRNA vs. Aβ+scr siRNA vs. Aβ+SHARPIN siRNA is 1: 0.96: 1.6, n=4 for TGF-β release) (**j, k**). Statistical analysis-One-way NOVA with *p>0.05, **p>0.01, ***p>0.001.

### 3.3. SHARPIN knockdown prevents inflammation-mediated neuronal cell death

The effect of SHARPIN-mediated inflammation on neuronal apoptosis was analyzed using conditioned media experiments. The differentiated SHSY5Y cells by treatment with 10 μM RA showed increased size and synaptic connections together with decreased expression of the neuronal stem cell marker, Nestin (Fig.4a).

**Fig.4:**
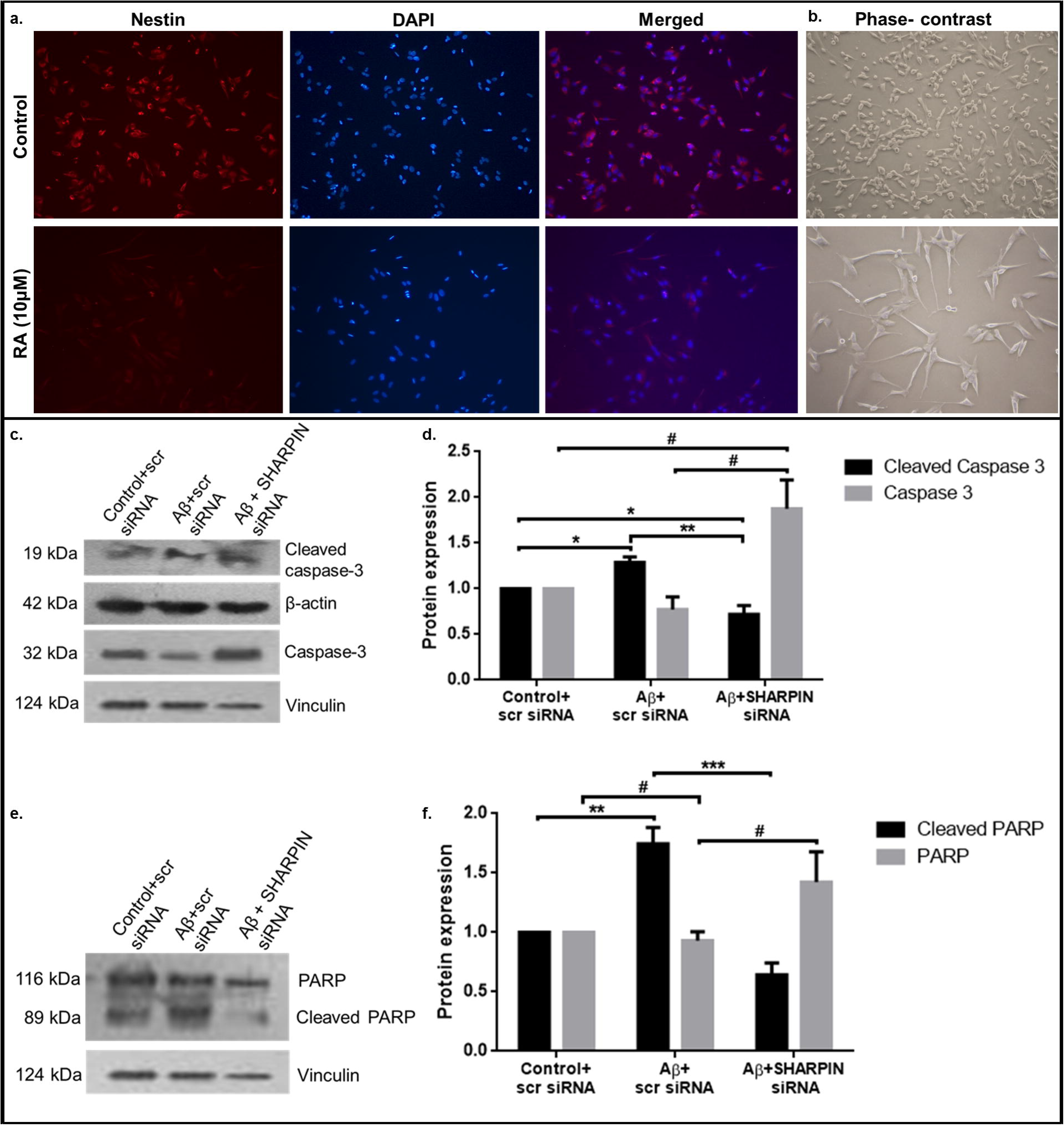
SHARPIN knockdown in macrophages prevents Aβ-induced inflammation mediated neuronal damage. SHSY5Y neuroblastoma cell lines were differentiated into mature neurons by incubating the cells with 10μM Retinoic acid (RA) for 3-4 days in RPMI-1640 supplemented with 1% FBS. The differentiated cells were analysed for the stem cell lineage marker, Nestin using immunocytochemistry and analysed using fluorescent imaging (**a**). The cells were further analysed for cell size, morphology and synaptic connections before and after differentiation using phase-contrast microscopy (**b**) (different field). Differentiated SHSY5Y cells were treated with conditioned media obtained from macrophages transfected with SHARPIN siRNA/scrambled siRNA and then treated with 10μM Aβ, for 24h. The expression of apoptotic markers cleaved caspase-3 (**c**) and cleaved PARP (**e**) using western blotting, normalised with control and represented graphically (Fold change in expression: Control+scr siRNA vs. Aβ+scr siRNA vs. Aβ+SHARPIN siRNA is 1: 1.29: 0.72, n=3 for cleaved caspase-3 and 1: 0.77: 1.88, n=3 for caspase-3; Control+scr siRNA vs. Aβ+scr siRNA vs. Aβ+SHARPIN siRNA is 1: 1.75: 0.64, n=3 for cleaved PARP and 1: 0.93: 1.42, n=3 for PARP) (**d, f**). Statistical analysis-One-way NOVA with *^/#^p>0.05, **p>0.01, ***p>0.001.

Neuronal cells treated with conditioned media obtained from macrophages incubated with Aβ showed increased expression of apoptotic markers cleaved caspase 3 and cleaved PARP. Importantly, incubation of the neuronal cells with conditioned media derived from SHARPIN-silenced THP-1 macrophages was found to significantly reduce the expression of pro-apoptotic markers (Fig.4).

### 3.4. Aβ-induced oxidative stress affects SHARPIN expression

Aβ is known to enhance Reactive Oxygen Species (ROS) generation and oxidative stress [18] in the AD brain and *in-vitro* cell cultures. In support of this findings in previous studies, we show enhanced ROS generation in Aβ-treated macrophages compared to the control. Addition of NAC in the presence of Aβ reduced ROS levels thereby providing evidence for the presence of Aβ stimulating ROS production in macrophages (Fig.5a, b). To check the role of ROS in stimulating SHARPIN expression in macrophages exposed to Aβ, the cells were pre-incubated with NAC and SHARPIN expression was analyzed. NAC treatment downregulated SHARPIN expression, demonstrating the role of ROS in mediating Aβ-stimulated SHARPIN expression in macrophages (Fig.5d, e).

**Fig.5:**
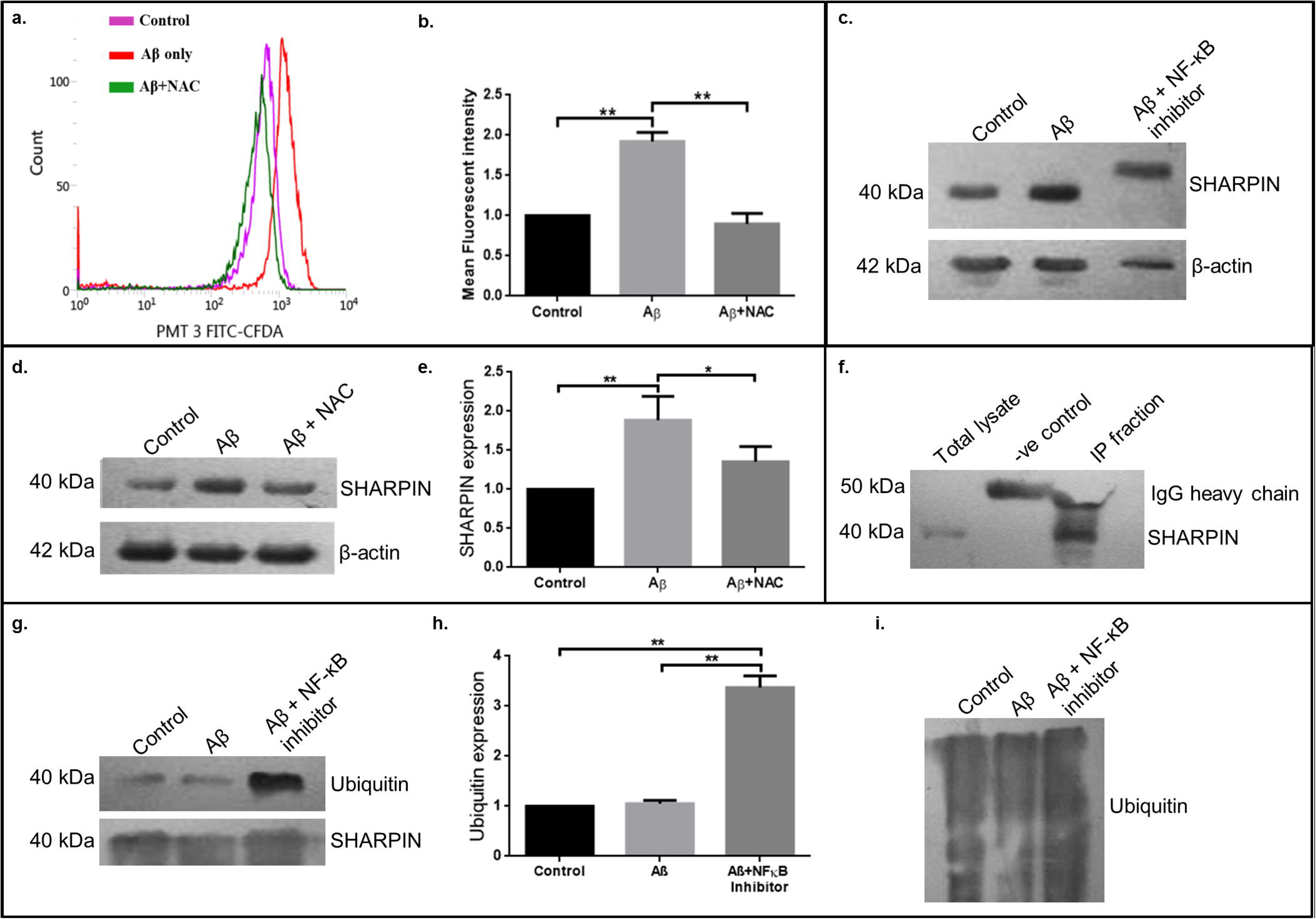
SHARPIN expression is stimulated by Aβ-induced oxidative stress and is post-translationally modified by NF-κB-mediated signalling pathway. Differentiated THP-1 macrophages were pre-incubated with 10mM NAC for 1h and then with NAC + 40μM Aβ for 12 h. Then the cells were incubated with H2DCFDA for 1h and subjected to flow cytometry for analysing intracellular reactive oxygen species (ROS) production induced by Aβ (**a**) and the mean fluorescent intensity was quantified, normalised with control and represented graphically (Fold change in expression: Control vs. Aβ vs. Aβ+NAC is 1: 1.92: 0.89, n=3) (**b**). Differentiated THP-1 macrophages were pre-incubated with 10mM NAC for 1h and then with NAC + 10μM Aβ for 6 h and the expression of SHARPIN was analysed using western blotting (**d**), normalised with control and was represented graphically (Fold change in expression: Control vs. Aβ vs. Aβ+NAC is 1: 1.89: 1.35, n=3) (**e**). **C.** Differentiated THP-1 macrophages were pre-incubated with NF-κB inhibitor, Bay 11-7082 (10μM) for 1h and then treated with Bay 11-7082 + 10μM Aβ for 12h. SHARPIN expression was analysed using western blotting and the increase in molecular weight of SHARPIN in Bay 11-7082 + 10μM Aβ treated cells compared to control and Aβ was represented (**c**). SHARPIN protein was immunoprecipitated for further analysis and the purity of the immunoprecipitated protein was shown with western blot data (**f**). Immunoprecipitated SHARPIN protein was probed for ubiquitination using anti-ubiquitin antibody and the ubiquitination of SHARPIN protein (**g**) and total protein ubiquitination (**i**) was analysed using western blotting, normalised with control and represented graphically (Fold change in expression: Control vs. Aβ vs. Aβ+ Bay 11-7082 is 1: 1.05: 3.37, n=3) (**h**). Statistical analysis-One-way NOVA with *p>0.05, **p>0.01, ***p>0.001.

### 3.5. SHARPIN is controlled by NF-κB-mediated feedback regulation

Since NF-κB is a redox-sensitive transcription factor, its regulatory role on SHARPIN was investigated. Pharmacological inhibition of NF-κB using Bay-117082 was found to cause an increase in the molecular weight of SHARPIN compared to the control and Aβ groups (Fig.5c). Immunoprecipitation of SHARPIN (Fig.5f) and probing with anti-ubiquitin antibody in the Aβ+Bay-117082 treated group showed a significant increase in ubiquitinylation (Fig.5g), suggesting that the transcription factor NF-κB may function to ubiquitinate the protein SHARPIN.

### 3.6. SHARPIN expression is altered in AD patient macrophages

Study subjects were categorized into 31 AD, 13 MCI and 19 control subjects (Fig.6a). We have observed that SHARPIN expression by macrophages was showing an increased expression in MCI subjects compared to AD and control subjects (Fig. 6b). Importantly, SHARPIN showed a positive trend in the expression patterns with Aβ_42/40_ release in the plasma of control, MCI and AD subjects (Fig.6d, e, f) which corresponds to the fact that SHARPIN expression by macrophages, in the absence of systemic inflammation, might be stimulated by the concentration of Aβ_42/40_ in the blood plasma. Due to the small sample size with a heterogeneous population, we tried to segregate the subjects on the basis of age (Fig.6g) and the presence of diabetes (Suppl. Fig.7S) and we could find an increased SHARPIN expression in subjects in a higher age group and with diabetes.

**Fig.6:**
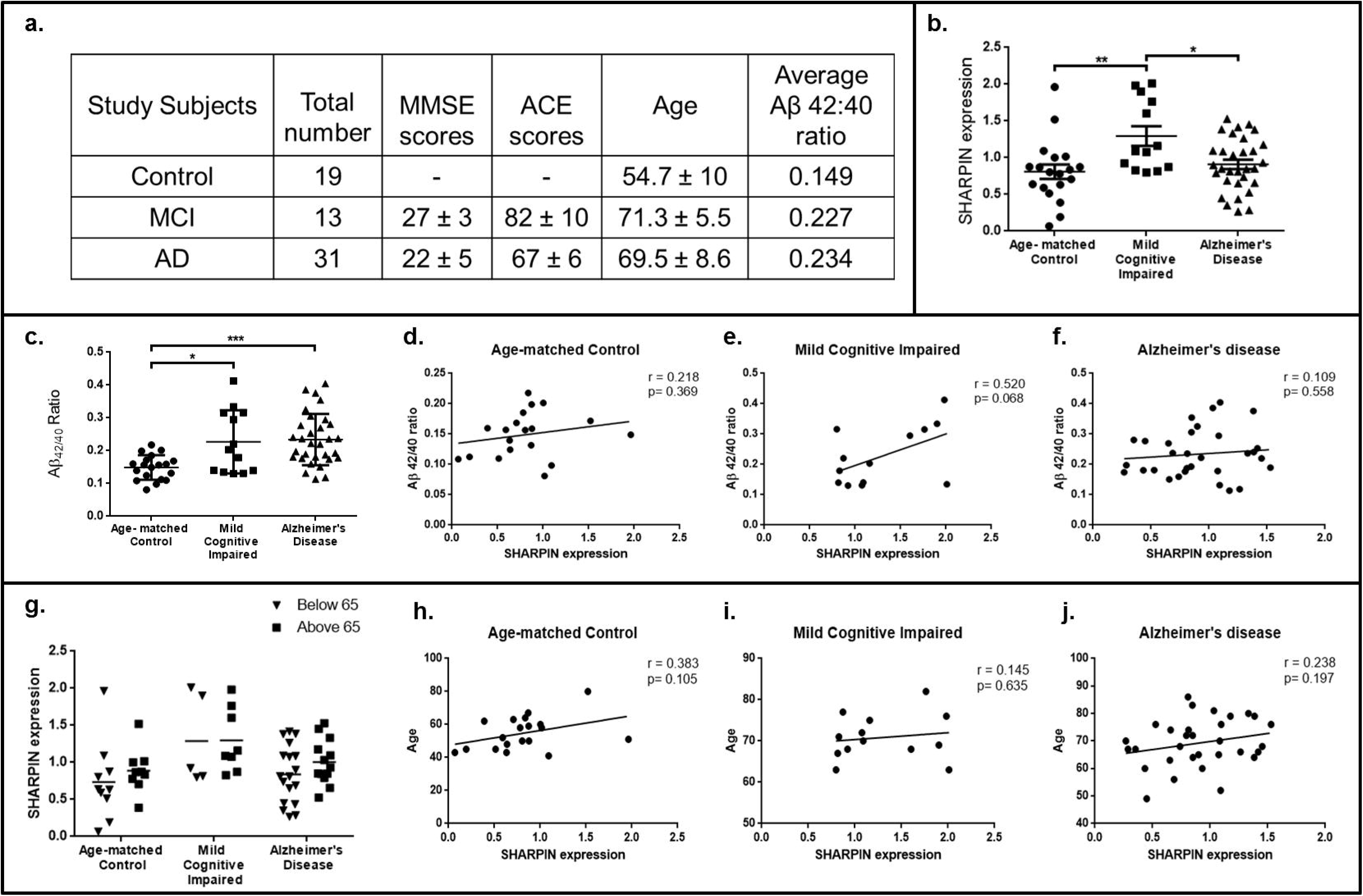
SHARPIN expression in Mild Cognitively Impaired (MCI), Alzheimer’s disease (AD) and control subjects. Details of the study subjects were represented. Subjects with high CRP level were excluded from the study population (**a**). Peripheral blood mononuclear cells (PBMCs) were isolated from the study subjects’ blood samples and were cultured in RPMI-1640 supplemented with autologous serum were cultured for 14 days until complete differentiation into macrophages. Total protein was isolated from the mature macrophages and analysed for SHARPIN expression in the study subjects and represented as scatter plot (**b**). Aβ_42_ and Aβ_40_ in the blood plasma isolated from the same study subjects were analysed using ELISA and the ratio of Aβ_42_ to Aβ_40_ was calculated and represented as scatter plot (**c**). SHARPIN expression by macrophages in the study subjects were correlated with Aβ_42/40_ ratio in the blood plasma of control (**d**), MCI (**e**) and AD (**f**) subjects. Pearson correlation coefficient, r = 0.218 (p=0.369, ns) for age-matched control, r = 0.520 (p=0.068, ns) for mild-cognitive impaired and r = 0.109 (p= 0.558 ns) for Alzheimer’s disease subjects (ns-non significant). The study subjects were grouped into age groups below 65 and above 65 and the expression of SHARPIN was analysed and represented as scatter plot (**g**). SHARPIN expression by the macrophages was correlated with age of the study subjects and represented graphically (**h, i, j**). Pearson correlation coefficient, r = 0.383 (p=0.105, ns) for age-matched control, r = 0.145 (p=0.635, ns) for mild-cognitive impaired and r = 0.238 (p= 0.197, ns) for Alzheimer’s disease subjects. *ns – non significant.

## 4. Discussion

Defective immune cell-mediated clearance of Aβ, is a major contributor of Aβ accumulation in the brain, leading to the pathogenesis of AD [19, 20]. Aβ accumulation-associated inflammatory activation of microglia in the brain and the macrophages entering the brain from peripheral circulation plays a major role in causing neuronal apoptosis, leading to amplified rate of progression of AD [21]. Although several studies have shown a correlation between inflammatory mediators and phagocytic receptor expression by immune cells [22], the underlying mechanisms that affect Aβ phagocytosis and promote pro-inflammatory conditions in the AD brain remain elusive.

SHARPIN has been recognised as an upstream activator of NF-κB thus acting as an important mediator of inflammatory mechanism [23–25]. A recent study by Yuya Asanomi et.al identified a rare functional variant of SHARPIN as a genetic risk factor for late-onset AD (LOAD) proving the role of the protein in the progression of AD [26]. While the study identified genetic mutations in SHARPIN as a risk factor for LOAD, the molecular basis of SHARPIN in AD pathogenesis remains unclear.

In the present study, we focused on the role of SHARPIN in the regulation of macrophage function and its contribution to the progression of AD. Using differentiated THP-1 macrophages exposed to Aβ as an *in-vitro* model, we have demonstrated a significant increase in the expression of SHARPIN in the presence of Aβ, indicating a link between Aβ exposure and SHARPIN expression in macrophages. Further, our study has shown the role of SHARPIN in regulation of Aβ phagocytosis through modulating the expression of receptors involved in Aβ uptake thus, affecting the overall cellular intake of Aβ which was evident by the significant reduction in FITC-Aβ fluorescence in SHARPIN-silenced cells.

SHARPIN and NLRP3 are widely regarded as the two principal mediators of inflammation. A study conducted in mice carrying mutant SHARPIN (*Sharpin*^*cpdm*^) that caused loss of SHARPIN function identified that SHARPIN is required for NLRP3 activation [12]. We found a reduction in NLRP3 expression in SHARPIN-knockdown macrophages exposed to Aβ demonstrating for the first time, a novel link between the two inflammatory mediators in the context of AD. Since SHARPIN regulates NLRP3 which is an activator of pro-inflammatory signalling, we analysed the expression of inflammatory markers in SHARPIN-silenced macrophages exposed to Aβ. We found that while Aβ polarizes macrophages to the M1 phenotype, knockdown of SHARPIN in the presence of Aβ prevented M1 polarization polarizing the cells to the neuro-protective M2 phenotype.

It has been reported that brain-resident and peripheral circulation-derived immune cells eliminate aggregated Aβ deposits in the brain, causing inflammatory response through the secretion of inflammatory cytokines and reactive oxygen species, leading to neuronal damage in AD. Since SHARPIN regulates inflammation that causes neuronal apoptosis, we sought to analyse the role of SHARPIN in mediating neuronal cell apoptosis. Using differentiated SHSY5Y neurons, we have observed significant reduction in the apoptosis of neurons treated with conditioned media derived from SHARPIN-knockdown macrophages compared to the cells treated with conditioned media from macrophages incubated with Aβ. While Aβ functioned through SHARPIN-mediated mechanisms to promote inflammatory cytokine release by the macrophages in the media that induced neuronal cell death, knockdown of SHARPIN in macrophages prevented the release of pro-inflammatory cytokines, leading to the survival of neurons in culture. The results clearly indicate that Aβ-induced neuronal apoptosis is primarily mediated through Aβ-mediated inflammatory mechanisms that involve SHARPIN. Thus our study proves SHARPIN as a critical protein that acts as a double-edged sword regulating phagocytosis and inflammation, where its downregulation reduces inflammation and protects inflammatory neuronal damage, but affecting immune-mediated phagocytosis and clearance of Aβ.

Since SHARPIN was found to regulate critical macrophage functions involving phagocytosis and macrophage polarization, the study explored the factors that could be involved in the regulation of Aβ -stimulated SHARPIN expression. Aβ is well known to induce oxidative stress through ROS production [18, 27]. Since Aβ induced the expression of SHARPIN, we explored the role of ROS in mediating Aβ -induced SHARPIN. Treatment of macrophages with NAC, a potent ROS scavenger, prevented Aβ -induced SHARPIN expression, demonstrating ROS to play an important role in mediating SHARPIN expression in response to Aβ. Since NF-κB is the redox-sensitive transcription factor, the role of NF-κB activation in regulating SHARPIN expression was explored and leads us to the finding that NF-κB promoted ubiquitination of SHARPIN. The functional implications and regulatory mechanisms of SHARPIN need to be explored in detail in both on an immunological basis and therapeutic basis. SHARPIN has been highly explored in many disease conditions, especially in cancer stressing out its multiple roles in many LUBAC-independent regulatory mechanisms [28, 29], suggesting that this modification that we have identified, if not targeted for a proteosomal degradation, might be inhibiting only the LUBAC-dependent SHARPIN function, promoting its other cellular functions, which is out of scope for this study.

Analysis of SHARPIN in AD and MCI patient-derived macrophages pointed out a significant increase in SHARPIN expression in MCI-derived macrophages compared to AD and control subjects. MCI is regarded as the preclinical stage of AD. It has been reported that with progression of AD, more of Aβ gets accumulated in the brain with reduced clearance to the peripheral system, resulting in a reduced concentration of Aβ in the blood plasma of AD subjects compared to the MCI subjects. The SHARPIN expression patterns in our study subjects can be reflected as a result of the stimuli induced by varied concentration of Aβ in the peripheral circulation, where a higher concentration of Aβ in MCI subjects induced the enhanced expression of SHARPIN by MCI macrophages and the lower concentration of Aβ in AD and control subjects contributes to a lower stimuli for SHARPIN expression. Hence, the reduced rate of Aβ clearance from the AD brain coupled with decreased Aβ concentration in peripheral circulation of AD patients may explain the decrease in SHARPIN expression in AD-patient-derived macrophages compared to MCI. A correlation study of SHARPIN expression with Aβ_42/40_ in the plasma of the same study subjects shows a positive trend suggesting the same. Due to the heterogeneity of the study population, we further classified the subjects on the basis of age and diabetes and found that SHARPIN expression was higher in a higher age group and those with diabetes pointing out the diversity of factors that could modulate SHARPIN expression and inflammatory condition *in-vivo*. Further, although we tried to correlate SHARPIN expression in macrophages derived from the study subjects with the phagocytic efficiency of macrophages and inflammatory markers in the plasma of the respective study subjects, a significant correlation was not found (Suppl. Fig.6S). This indicates the presence of complex mechanisms regulating inflammation and phagocytosis under *in-vivo* conditions contrary to the controlled *in-vitro* setting. An elaborate study on a larger cohort in a lesser heterogenic population needs to be conducted to explore the role of SHARPIN in the peripheral system *in-vivo*.

In summary, our study demonstrates, for the first time, a novel role for SHARPIN in the regulation of macrophage function in response to Aβ in a setting of AD. SHARPIN was found to regulate Aβ phagocytosis and Aβ-stimulated inflammatory mechanisms in macrophages and we found that Aβ-induced SHARPIN-mediated inflammation triggered neuronal cell death, demonstrating its role in promoting neurodegeneration in AD. Oxidative stress, with its calamitous role in age-associated diseases, was found to be regulating the protein expression, which was further modulated by NF-κB-mediated post-translational modification. Importantly, SHARPIN expression varied with the levels of Aβ_42/40_ in patient-derived macrophages demonstrating a link between SHARPIN expression and AD progression. Future studies need to address the role of SHARPIN in AD using a larger cohort of study subjects in order to establish the role of this protein in AD pathogenesis. Further, the functional role of ubiquitination in altering SHARPIN function and its implications in macrophages needs to be addressed. Importantly, exploring microglial- and astrocyte-mediated SHARPIN expression and its role in phagocytosis and inflammatory pathways are relevant in this field since these are the cells that respond primarily to Aβ accumulation in AD brain.

## Acknowledgements

We thank all the patients and healthy volunteers involved in the study.

We thank Dr. Lakshmi. S, Additional Professor, Molecular medicine, Division of Cancer Research, Regional Cancer Centre, Trivandrum, for providing valuable suggestions regarding flow cytometry analysis. Further, we thank CSIR-NIIST, Trivandrum for kindly providing SHSY5Y cell line for conducting a part of our study.

## Abbreviations

AD: Alzheimer’s disease
Aβ: amyloid-beta
MCI: Mild Cognitive Impairment
SHARPIN: Shank-associated RH domain-interacting protein
NLRP3: nucleotide-binding domain (NOD)-like receptor protein 3
ASC: Apoptosis-associated Speck-like protein containing a caspase-recruitment domain (CARD)
LUBAC: linear ubiquitination assembly complex
iNOS: induced Nitric Oxide Synthase
IL-1β: Interleukin-1beta
TGF-β: Transforming Growth Factor-1beta
TNF-α: Tumor Necrosis Factor-alpha
PBMC: Peripheral Blood Mononuclear cells
NF-κB: Nuclear Factor kappa-light chain-enhancer of activated B cells
ROS: reactive oxygen species
CRP: C-Reactive Protein
PBS: Phosphate-buffered saline
RPMI: Roswell Park Memorial Institute
FBS: fetal bovine serum
THP-1: Tohoku Hospital Pediatrics-1
NH_4_OH: Ammonium hydroxide
FITC: Fluorescein isothiocyanate

## Author Contributions

SG, MPS, RNM and DK conceived and designed the study. DK performed the experiments and acquired the results. DK, MPS, RNM and SG analysed, interpreted the results and drafted the manuscript. All authors reviewed and approved the manuscript.

## Submission declaration and verification

All authors verify that the data contained in the manuscript being submitted have not been published previously (except in the form of an abstract, a published lecture or academic thesis, and preprint version in BioRxiv), that it is not under consideration for publication elsewhere, that its publication is approved by all authors and tacitly or explicitly by the responsible authorities where the work was carried out, and that, if accepted, it will not be published elsewhere in the same form, in English or in any other language, including electronically without the written consent of the copyright-holder.

## Compliance with Ethical Standards

### Ethical Approval

All procedures performed in the above study were in accordance with the ethical standards of the Institutional Human Ethical Committee (IEC/234/2009) and with The Code of Ethics of the World Medical Association (Declaration of Helsinki), 1964 and its later amendments or comparable standards.

### Informed Consent

Informed consent was obtained from all individual participants included in the study.

