## Supplementary figures and images for "A novel role for SHARPIN in amyloid-β phagocytosis and inflammation by peripheral blood-derived macrophages in Alzheimer’s disease"

### Supplementary figure 1S

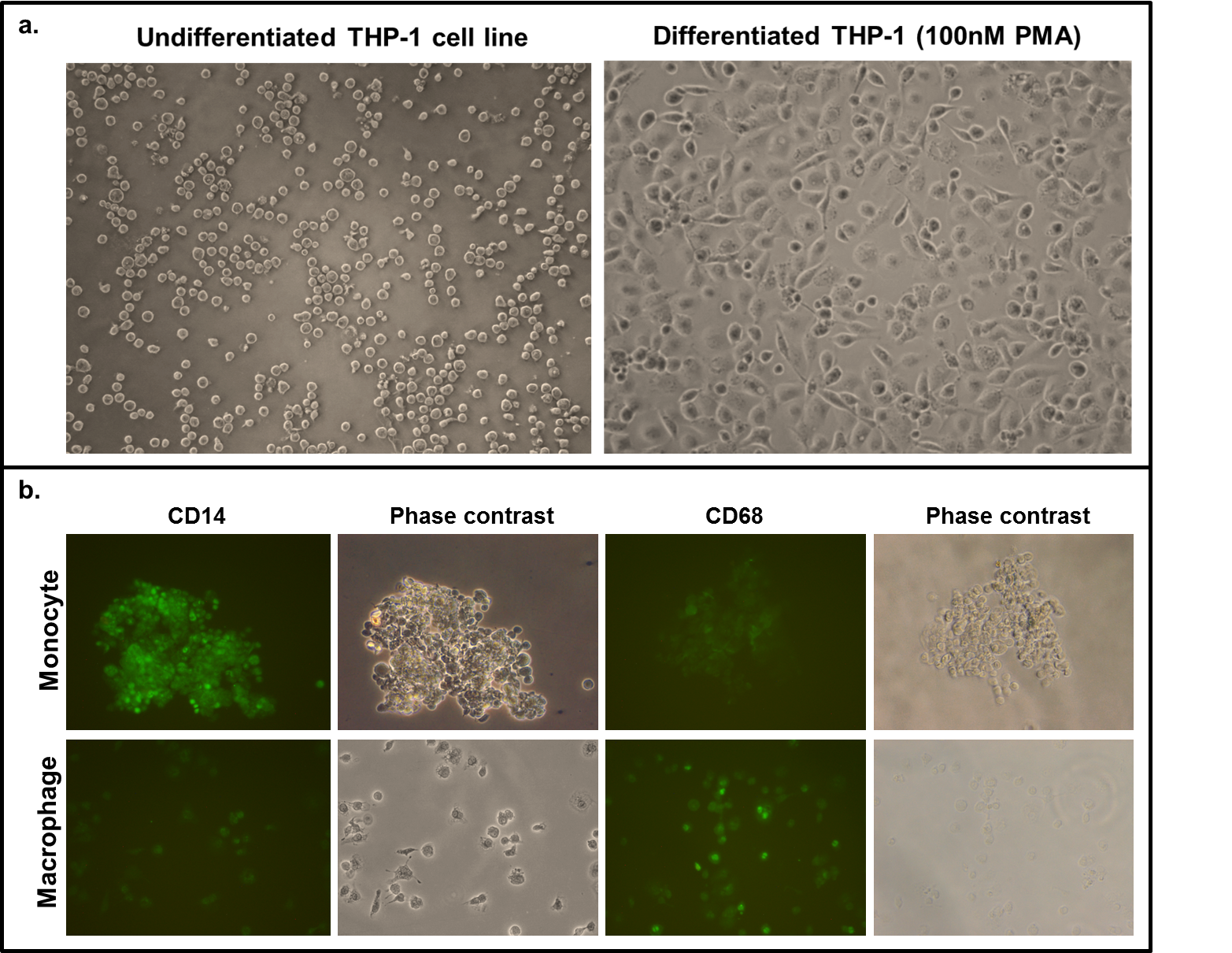

### Supplementary figure 3S

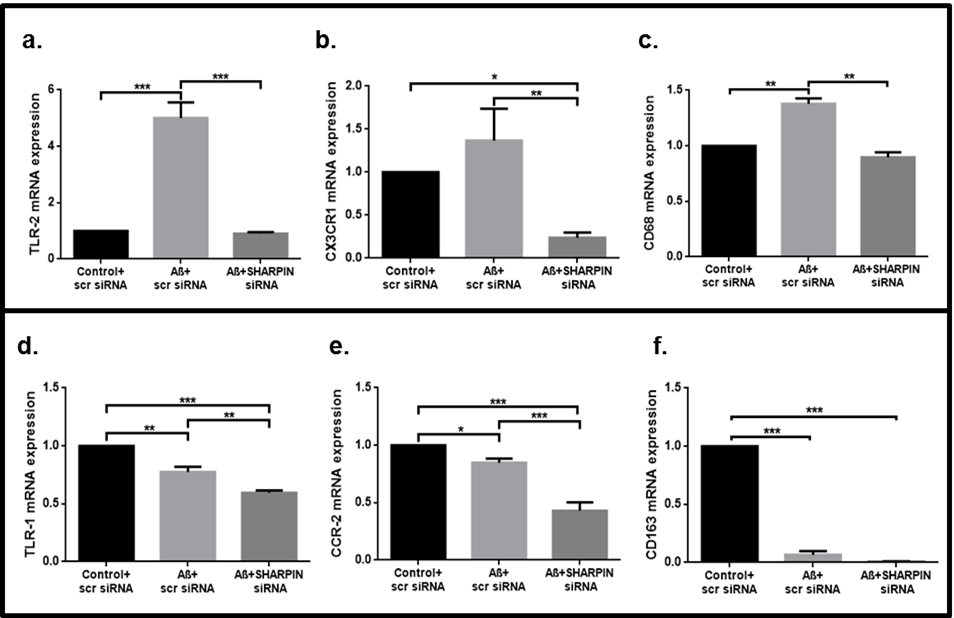

### Supplementary figure 6S

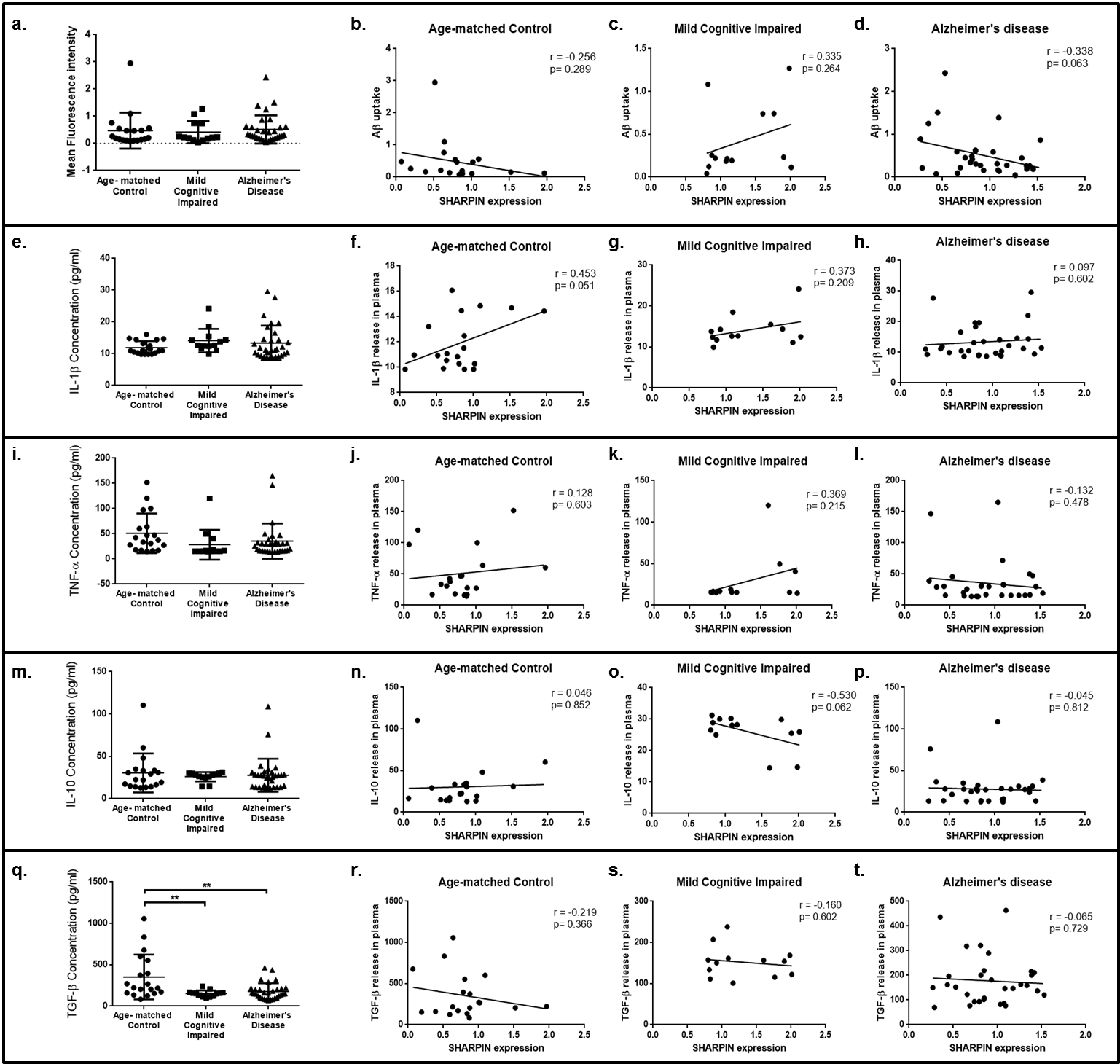

### Supplementary Figure 7S

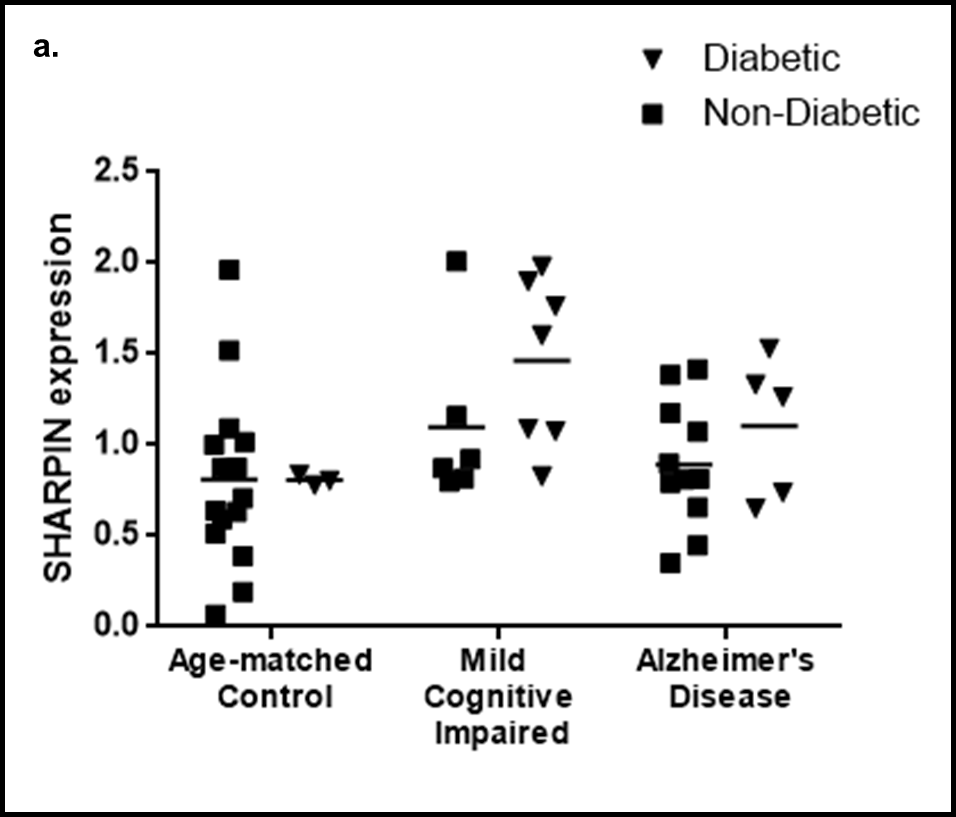
