## Supplementary figure legends for "A novel role for SHARPIN in amyloid-β phagocytosis and inflammation by peripheral blood-derived macrophages in Alzheimer’s disease"

**Supplementary Fig. 1S: Differentiation of THP-1 monocytes into macrophages.** THP-1 acute monocyte leukaemia cells were treated with 100nM phorbol 12-myristate 13-acetate (PMA) for 48 h in RPMI-1640 supplemented with 10% FBS for differentiation of monocytes into mature macrophages and the phase contrast image of cells before and after differentiation was analysed (**a**). THP-1 cells before and after PMA- induced differentiation were analysed for monocyte and macrophage markers CD14 and CD68 respectively using immunocytochemistry and analysed using fluorescent imaging (**b**).

**Supplementary Fig. 3S: SHARPIN knockdown inhibits Aβ-mediated M1 polarization of macrophages.** Differentiated THP-1 cells transfected with SHARPIN/scrambled (scr) siRNA were treated with 10μM Aβ for 12 h and the mRNA expression of M1 markers; TLR-2 (**a**), CX3CR1 (**b**) and CD68 (**c**) and M2 markers; TLR-1 (**d**), CCR2 (**e**) and CD163 (**f**) were analysed using Real-time PCR with tubulin as endogenous control and represented graphically after normalising with control. However, the mRNA expression of M2 markers also got decreased possibly because SHARPIN mediated signaling mechanisms are activating NF-κB and inhibition of SHARPIN thus downregulates the transcription of genes controlled by NF-κB which includes the M2 markers. Statistical analysis- One-way NOVA with *$p>0.05$, **$p>0.01$, ***$p>0.001$.

**Supplementary. Fig.6S: Correlation of SHARPIN expression by macrophages with phagocytosis and inflammation in study subjects.** Peripheral blood mononuclear cells (PBMCs) were isolated from the study subjects' blood samples and were cultured in RPMI-1640 supplemented with autologous serum were cultured for 14 days until complete differentiation into macrophages. Aβ phagocytic efficiency of the macrophages were analysed at the 14th day by incubating the cells with FITC-labelled Aβ overnight and the mean fluorescent intensity per cell (MFI) was calculated over three different fields per sample and represented as scatter plot (**a**).  MFI of each study sample was correlated with SHARPIN expression by the macrophages of the same subjects. Pearson correlation coefficient, r = -0.256 (p=0.289, ns) for age-matched control (**b**), r = -0.335 (p=0.264, ns) for mild-cognitive impaired (**c**) and r = -0.338 (p= 0.063, ns) for Alzheimer's disease (**d**) subjects. The release of pro-inflammatory cytokines IL-1β (**e**), TNF-α (**i**) and anti-inflammatory cytokines IL-10 (**m**), TGF-β (**q**) in the blood plasma of the study subjects were analysed using ELISA and represented graphically. SHARPIN expression by macrophages in the study subjects were correlated with cytokine release in the blood plasma. Pearson correlation coefficient for IL-1β, r = 0.453 (p=0.051, ns) for age-matched control (**f**), r = 0.373 (p=0.209, ns) for mild-cognitive impaired (**g**) and r = 0.097 (p=0.602, ns) for Alzheimer's disease (**h**) subjects. Pearson correlation coefficient for TNF-α, r = 0.128 (p=0.603, ns) for age-matched control (**j**), r = 0.369 (p=0.215, ns) for mild-cognitive impaired (**k**) and r = -0.132 (p=0.478, ns) for Alzheimer's disease (**l**) subjects. Pearson correlation coefficient for IL-10, r = 0.046 (p=0.852, ns) for age-matched control (**n**), r = -0.530 (p=0.062, ns) for mild-cognitive impaired (**o**) and r = -0.045 (p=0.812, ns) for Alzheimer's disease (**p**) subjects. Pearson correlation coefficient for TGF-β, r = -0.219 (p=0.366, ns) for age-matched control (**r**) , r = -0.160 (p=0.602, ns) for mild-cognitive impaired (**s**) and r = -0.065 (p= 0.729, ns) for Alzheimer's disease (**t**) subjects (ns- non significant). *ns – non significant.

**Supplementary Fig. 7S: SHARPIN expression in study subjects categorized as diabetic and non-diabetic subjects.** The study subjects were categorised on the basis of presence of diabetes and the expression of SHARPIN by the macrophages of the subjects were analysed using western blotting and represented graphically (**a**).
